# Unveiling multi-scale architectural features in single-cell Hi-C data using scCAFE

**DOI:** 10.1101/2024.09.10.611762

**Authors:** Fuzhou Wang, Jiecong Lin, Hamid Alinejad-Rokny, Wenjing Ma, Lingkuan Meng, Lei Huang, Jixiang Yu, Nanjun Chen, Yuchen Wang, Zhongyu Yao, Weidun Xie, Xiangtao Li, Ka-Chun Wong

## Abstract

Single-cell Hi-C (scHi-C) has provided unprecedented insights into the heterogeneity of 3D genome organization. However, its sparse and noisy nature poses challenges for computational analyses, such as chromatin architectural feature identification. Here, we introduce scCAFE, a deep learning model for the multi-scale detection of architectural features at the single-cell level. scCAFE provides a unified framework for annotating chromatin loops, TAD-like domains (TLDs), and compartments across individual cells. Our model outperforms previous scHi-C loop calling methods and delivers accurate predictions of TLDs and compartments that are biologically consistent with previous studies. The resulting single-cell annotations also offer a measure to characterize the heterogeneity of different levels of architectural features across cell types. We leverage this heterogeneity and identify a series of marker loop anchors, which demonstrate the potential of the 3D genome data to annotate cell identities without the aid of simultaneously sequenced omics data. Overall, scCAFE not only serves as a useful tool for analyzing single-cell genomic architecture, but also paves the way for precise cell-type annotations solely based on 3D genome features.

## Introduction

Chromatin architecture plays a crucial role in gene regulation and cellular function. High-throughput chromosome conformation capture (Hi-C) (1, 2) has significantly enhanced our understanding of three-dimensional (3D) genome organization by providing comprehensive maps of chromatin interactions throughout the entire genome. Although the molecular basis and driving mechanisms of 3D chromatin folding are not yet fully understood, extensive research has reached a consensus that the genome is organized hierarchically in 3D space (3–5). This hierarchical organization can be broadly categorized into three layers: compartments, topologically associating domains (TADs), and chromatin loops. These layers of organizational units are ubiquitous features present throughout the whole genome (4, 6) and have been revealed to perform critical functions in various biological processes, such as DNA replication (7) and gene regulation (8, 9). The understanding on the multiple levels of genome architecture largely comes from the population-level mapping of genomic loci proximity using Hi-C-like techniques (e.g., bulk Hi-C). From the contact maps obtained from these assays, the architectural features exhibit distinct patterns and therefore can be easily identified using computational methods (6, 10).

However, the patterns observed from these experiments only reflect an ensemble of the genome organization in all the individual cells. This means that the data represents an average view, masking the heterogeneity and unique configurations present across individual cells. To overcome this limitation, single-cell Hi-C (scHi-C) has emerged as a powerful technique, facilitating the examination of genome architecture at the single-cell level (11–17). scHi-C data offers unparalleled insights into the heterogeneity of chromatin organization among individual cells, unveiling distinct structural configurations that are typically obscured in bulk analyses. This single-cell lens is especially vital for understanding the dynamic nature of chromatin interactions in various biological contexts (18), including development (19), differentiation (20), and aging throughout the lifespan (17). However, the sparse and noisy characteristics of scHi-C data present substantial challenges for computational analysis, that the visual patterns typically observed in bulk Hi-C contact maps are significantly diminished in scHi-C data. This has necessitated the development of robust methods specifically designed to detect architectural features in scHi-C data.

Multiple previous studies have converged on a paradigm for scHi-C architectural feature analysis, which can be summarized as an “imputation-and-calling” strategy. First, the contact maps are imputed to a density equivalent to bulk levels to recover the visual patterns of the architectural features. Then, algorithms that operate on the dense contact maps are applied (21–27). Despite the remarkable performance these methods offer, this strategy often demands substantial computational resources or extended execution time. Moreover, the reliance on imputation may introduce biases and limit the achievable resolution, potentially obscuring fine-grained architectural features. To facilitate a more efficient identification of architectural features at the single-cell level, several tools have been published recently for single-cell loop detection (28) and single-cell TAD-like-domain (TLD) identification (29, 30). However, to the best of our knowledge, there has not been an integrative framework capable of pre-dicting all three levels of 3D genome features at the singlecell level without relying on imputation. This absence can increase the complexity and difficulty of practical applications. Additionally, an integrated framework for multi-layer prediction could provide a unified understanding of the dynamics of architectural features, offering a comprehensive view of spatial genome organization. With the multi-scale representation extracted, such an approach has the potential to reveal novel insights into the intricate relationship between 3D genome features and cellular functions.

In this study, we introduce scCAFE (Calling Architectural FeaturEs at the single-cell level), a comprehensive framework that utilizes multi-task learning techniques to predict 3D architectural elements from scHi-C data without relying on dense imputation. scCAFE is designed to simultaneously predict chromatin loops and reconstruct sparse contact maps, thereby generating a highly expressive representation of genome organization. Following the acquisition of these embeddings, we apply unsupervised learning methods to identify TLDs and compartments. scCAFE surpasses existing state-of-the-art methods in architectural feature calling. Leveraging the feature calls generated by scCAFE, we assess the predictive capabilities of various layers of single-cell architectural features in determining cell identity. Our findings reveal that compartments and loops serve as better predictors of cell identity compared to TLDs. Additionally, we investigate the feasibility of identifying cell types based solely on loop frequencies of each loop anchor. These insights underscore the potential of scCAFE in advancing our understanding of genome organization and its implications for cellular differentiation and identity.

## Results

### Constructing machine learning models to detect single-cell 3D architectural features

To detect 3D architectural features from the extremely sparse single-cell Hi-C data without densely imputing the contact maps, we designed a neural network with multi-task learning techniques. We illustrated the overall architecture of scCAFE in Figure 1. The multi-task variational graph autoencoder (VGAE) proposed in this study was trained to simultaneously predict chromatin loops and reconstruct contact maps. This approach enabled the model to learn latent features that effectively captured both loop patterns and the global 3D organizational features of chromosomes. The representations learned via this method were beneficial for predicting chromatin loops and enabled the unsupervised identification of TLDs and compartments at the single-cell level.

**Fig. 1.**
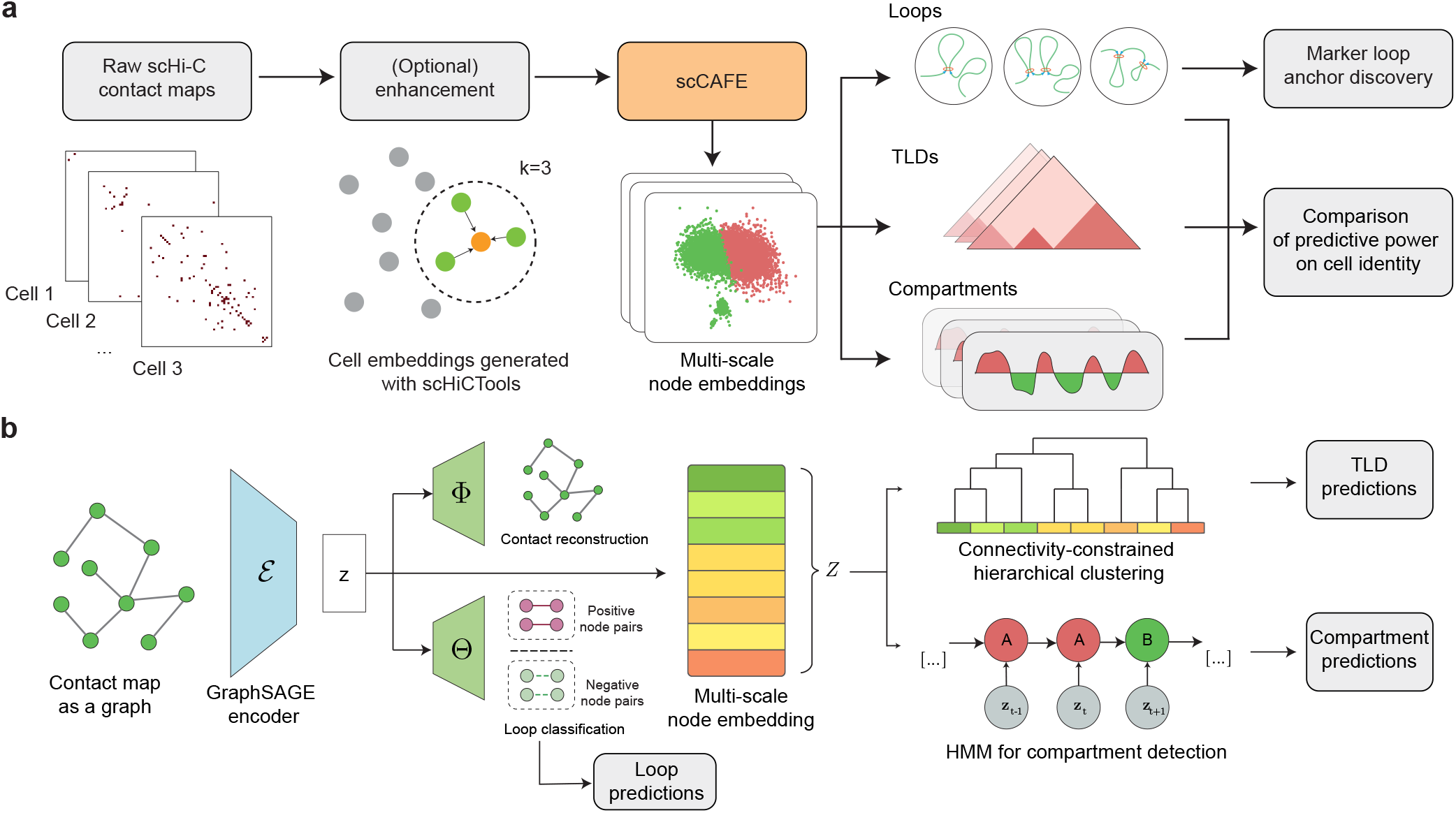
Overview of the scCAFE framework. **(a)** Workflow of scCAFE. **(b)** Model architecture of scCAFE. Each input contact map is treated as a graph and passed through a GraphSAGE encoder to generate latent variables. These latent features are then decoded by two decoders, Φ and Θ, to reconstruct the original contact maps and classify the loops, respectively. Subsequently, the latent features are treated as an ordered sequence. They are input to a connectivity-constrained hierarchical clustering model for TLD predictions and fed to a hidden Markov model (HMM) for compartment predictions.

In the framework of scCAFE, the chromosome graphs were not densified to recover the data resolution of bulk Hi-C data. Instead, only graphs of poor quality were slightly enhanced using an enhancement module. Despite this enhancement, the contact maps remained sparse, maintaining the computational efficiency of the model. The model’s predictive efficacy for multi-scale organizational units (introduced in later sections) also validated that 3D architectural features could be directly inferred from the sparse contact matrices without the need for dense imputation.

An important principle we followed in constructing the model was the multi-view nature of Hi-C data (28, 31). Consistent with scGSLoop (28), in scCAFE, the neural network operates on the graph view generated from contact maps, with node (genomic bin) features derived from DNA sequences, reflecting the sequence view. Moreover, we leveraged the sequence characteristics of the data to predict higher-order organizational units at the single-cell level, including TLDs and compartments. Given that the genomic bins have an order on the linear genome, the node embeddings of a chromosome generated by the neural network can be considered as a linearly connected sequence, which allowed us to apply sequence-based learning algorithms effectively. In particular, hierarchical clustering with a linear connectivity constraint was adopted for TLD calling, while an HMM was utilized for compartment identification. The detailed approach is outlined later in the Methods section.

A simple yet effective method for marker loop anchor identification was proposed and integrated into scCAFE. By utilizing chromatin loops detected at the single-cell level, scCAFE recognizes 10 kb-genomic bins that are significantly enriched with chromatin loops in different cell types. Except for single-cell predictions of 3D genome features at different scales, the scCAFE model also provides an interface to predict these features at the consensus level. This dual capability enables researchers to investigate the spatial structures of the genome both within individual cells and across cell populations, offering a comprehensive view of 3D genomic organization.

### scCAFE predicts loops accurately for both single cells and cell populations

We conducted a comprehensive assessment of scCAFE’s performance in predicting chromatin loops at both the single-cell and consensus levels. The datasets that we used for performance evaluation include the mouse embryonic cells (mESC) dataset from (12) and the human prefrontal cortex (hPFC) dataset from (32). First, we compared our method against scGSLoop (28) to evaluate its ability in identifying loops at the single-cell level. The F1 score, precision, and recall were calculated for each individual cell using the single-cell loop calls. The results of the comparisons in terms of these metrics are presented in Figure 2a-d, illustrating the effectiveness of the model in accurately predicting single-cell loops. By conducting Wilcoxon signed-rank tests on the F1 scores, precisions, and recalls of scGSLoop and scCAFE, it was shown that scCAFE significantly outperformed scGSLoop in all the cell types of the two datasets (denoted by the asterisks in Figure 2a-d).

**Fig. 2.**
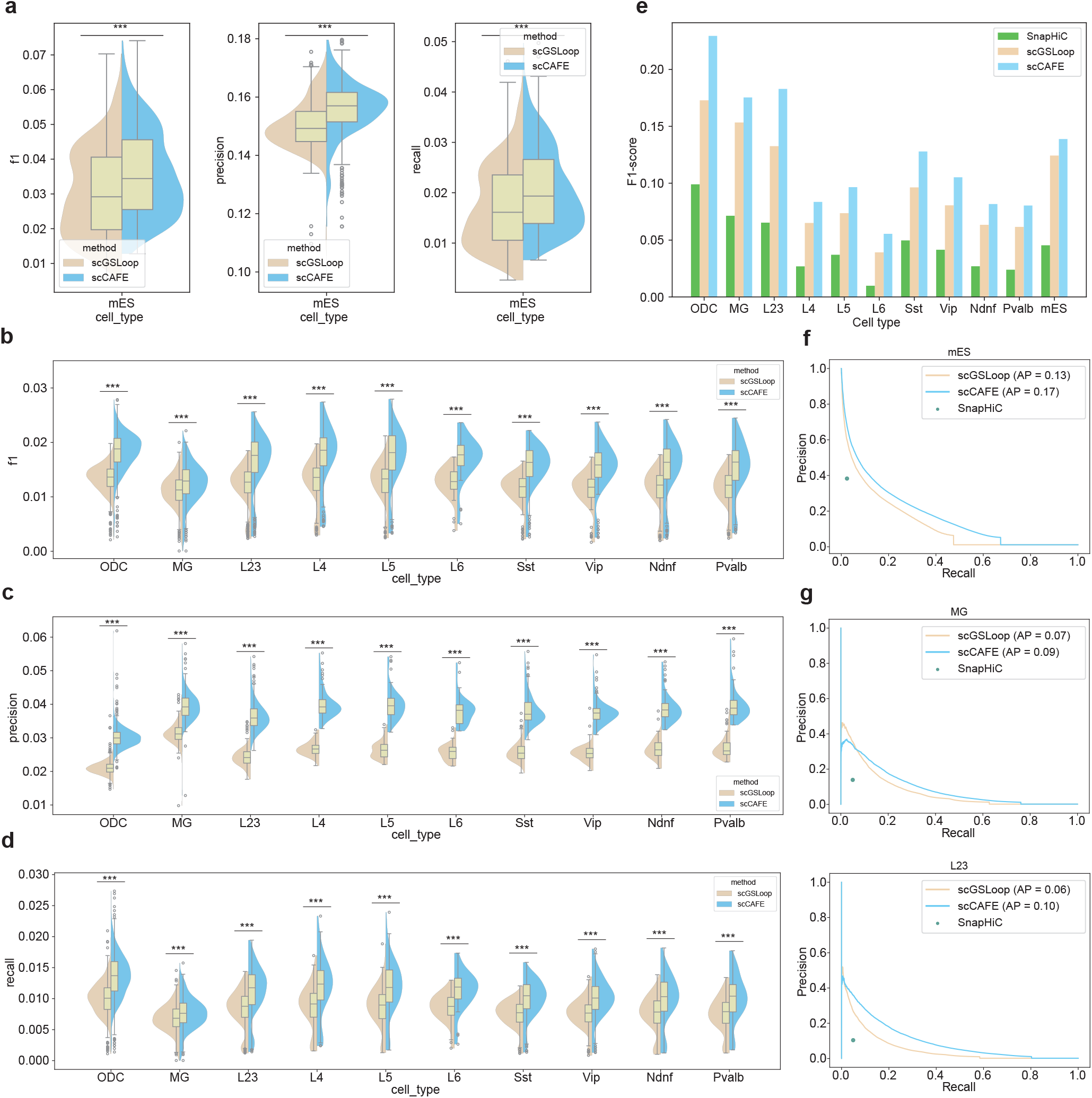
scCAFE accurately detects chromatin loops at both the single-cell level and the consensus level. **(a)** F1 scores, precisions, and recalls of single-cell chromatin loops in the mESC dataset predicted by scCAFE and scGSLoop. Three asterisks (***) indicate a p-value smaller than 0.001, determined using the Wilcoxon signed-rank test (the same in panels b-d). **(b-d)** F1 scores, precisions, and recalls of single-cell loops in different cell types of the hPFC dataset, compared between scCAFE and scGSLoop. **(e)** F1 scores for consensus loops across different cell types in both datasets, comparing the performance of SnapHiC, scGSLoop, and scCAFE. **(f)** Precision-recall plots of the consensus loops in the mESC dataset predicted by SnapHiC, scGSLoop, and scCAFE, respectively. **(g)** Precision-recall plots of the consensus loops in MG and L2/3 cells predicted by SnapHiC, scGSLoop, and scCAFE, respectively. The precision-recall curves in other cell types of the hPFC dataset can be found in Supplementary Figure S1.

In certain studies, researchers focus particularly on the chromatin loops of various cell types and sub-types. Consequently, it is crucial that a single-cell loop-calling method is capable of predicting chromatin loops summarizing the average looping patterns of the cell population. In scCAFE, we adopted the same aggregation method as in scGSLoop to predict consensus loops (i.e., the loops at the population level). It was demonstrated that the consensus loop predictions consistently achieved higher F1 scores compared to SnapHiC and scGSLoop across all tested cell types (Figure 2e). Additionally, in order to comprehensively examine the performance of scCAFE in terms of consensus loop prediction, we conducted additional evaluations across various probability thresholds using PR curves. As shown in Figure 2f,g and Supplementary Figure S1, the PR curves of scCAFE exhibited higher precision across various recall values compared to the scGSLoop model in 9 out of 11 cell types from both the mESC and hPFC datasets. In both datasets, scCAFE achieved higher average precision (AP) than scGSLoop across all cell types. Additionally, the points representing SnapHiC predictions were consistently positioned lower than the PR curves of both scGSLoop and scCAFE. These results suggest that the algorithmic design of our method improves loop predictions at the consensus level, providing better performance than previous approaches.

In all the performance assessments described above, we adhered to the principle of training the model on one dataset and testing it on a different dataset. This approach ensures a fair comparison and better simulates real-world applications. The remarkable performance, despite differences in data conditions and species, highlights the model’s ability to generalize across varying scenarios.

### Detection of TLDs in scHi-C data using scCAFE

scCAFE has empowered us to unveil multiple layers of 3D organizational elements within the genome through the utilization of unsupervised learning algorithms. In addition to identifying chromatin loops, we have successfully detected higher-order elements, such as TLDs and compartments, at both the single-cell and consensus levels. This section focuses on evaluating the performance of scCAFE in TLD calling. Specifically, we identified single-cell TLDs and consensus TLDs in the mESC dataset. Figure 3a illustrates an example region of the mESC genome, highlighting the TLDs identified by scCAFE. It is shown that the predictions generated by scCAFE at both the single-cell level and the consensus level were highly consistent with the patterns observed in bulk Hi-C. In particular, the single-cell TLD and consensus TLD boundaries aligned with the local minima of insulation scores and previously reported boundary-enriched factors, such as CTCF, H3K4me3, and H3K27ac (33, 34).

**Fig. 3.**
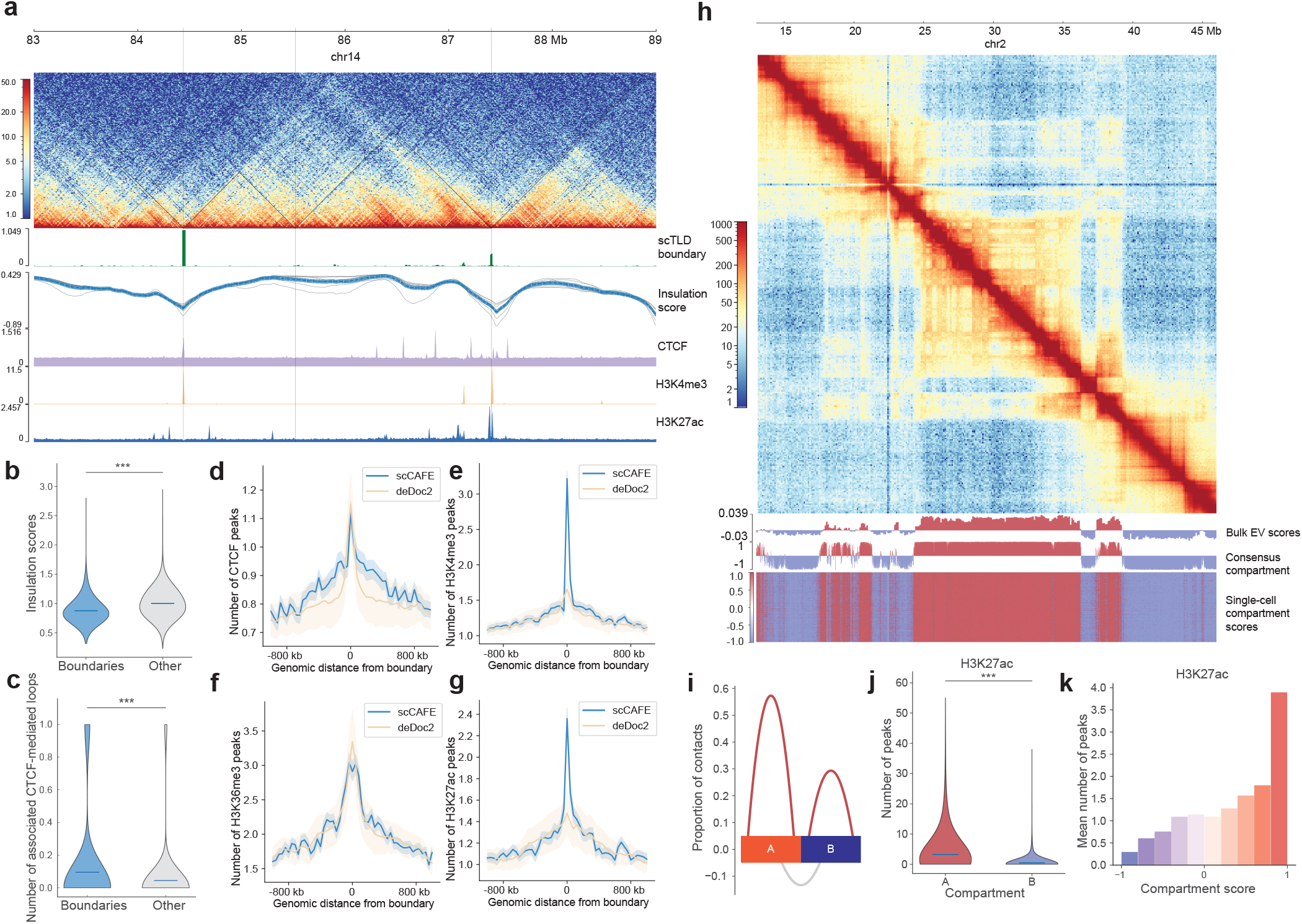
TLDs and compartments predicted by scCAFE are validated by bulk Hi-C and epigenetic features. **(a)** An example region in the genome of mESC, annotated with scCAFE-predicted consensus TLDs and single-cell TLDs. The insulation scores detected in bulk Hi-C data, as well as the tracks of CTCF, H3K4me3, and H3K27ac ChIP-Seq are also annotated. The predicted TLDs show consistency with the bulk insulation scores and epigenetic features. **(b)** The bulk insulation scores at the boundary regions and non-boundary regions. Three asterisks denote significance level of p-value smaller than 0.001, Mann–Whitney U test (same for panel c). **(c)** The numbers of CTCF-mediated loops detected at boundary regions and non-boundary regions by ChIA-PET. **(d-g)** Enrichment profiles of CTCF, H3K4me3, H3K36me3, and H3K27ac ChIP-Seq peaks around single-cell TLD boundaries detected by scCAFE and deDoc2. Shaded areas represent +/standard deviation. **(h)** An example genomic region showing scCAFE-predicted single-cell and consensus compartment scores. **(i)** Bulk Hi-C interactions within and between compartments. **(j)** Numbers of H3K27ac ChIP-Seq peaks in compartment A and B. **(k)** Numbers of H3K27ac peaks at different scCAFE compartment scores.

We conducted a series of genome-wide analyses to investigate the structural and functional characteristics of the identified TLDs. First, we compared the insulation scores between the boundary regions and the non-boundary regions. Previous studies have shown a strong correlation between TAD boundary occurrences and lower insulation scores (35). In our analysis, the lower insulation scores derived from bulk Hi-C data serve as a validation marker for the consensus TLD boundaries. As illustrated in Figure 3b, the insulation scores are significantly lower at the consensus boundary regions detected by scCAFE, indicating a clear demarcation of TLD boundaries.

We next investigated the enrichment of CTCF and histone modifications at these boundaries as detected by scCAFE.

Figure 3d-g demonstrate a clear enrichment of CTCF binding, H3K4me3, H3K36me3, and H3K27ac at the single-cell TLD boundaries detected by our method. This finding aligns with the established enrichment profiles reported in previous research (33). We also compared these enrichment profiles with the ones detected using the state-of-the-art single-cell TLD calling method deDoc2 (30). It is shown in Figure 3d-g that the enrichment of CTCF and H3K36me3 of scCAFEpredicted TLD boundaries is slightly less pronounced than deDoc2 predictions. However, our model produced much more enriched profiles of H3K4me3 and H3K27ac, indicating a distinct, functionality-related TLD pattern predicted by scCAFE. Another layer of relevant structural properties of TLD boundaries is that they usually co-exist with CTCF-mediated loops (2, 3, 36). Hence, we utilized an independent ChIA-PET dataset on CTCF (37) to validate our model’s pre-dicted TLD boundaries. By integrating this dataset, we were able to confirm that our consensus TLD boundaries are associated with significantly higher numbers of CTCF-mediated loops compared with non-boundary regions (Figure 3c).

Taken together, the TLDs detected by scCAFE exhibit a comprehensive range of typical organizational and functional characteristics as recognized in previous studies. The enrichment comparison between our model and the state-of-the-art single-cell TLD calling method has also highlighted its remarkable performance. These findings underscore the ability of scCAFE in identifying TLDs in scHi-C data.

### Identification of A/B compartments in single cells using scCAFE

We further applied an HMM on the embeddings generated by scCAFE to identify A/B compartments in single cells, which represent an architectural feature at the higher level of the 3D genome hierarchy. Using this method, we identified the single-cell compartments and consensus compartments in the mESC dataset. Figure 3h presents an ex-ample region in the mESC genome where the checkerboard pattern can be clearly observed. By plotting the eigenvectors of the correlation matrix of bulk Hi-C data alongside our predicted single-cell and consensus compartments, we observed that the compartment patterns identified by scCAFE strongly coincide with the bulk eigenvectors and the patterns on the contact map.

By investigating the organizational and functional properties of the A/B compartments detected with our model, we aimed to further validate the compartment annotations produced by scCAFE. First, we elucidated the volume of chromatin contacts (from bulk Hi-C data) within compartments A and B, as well as between different compartments. As shown in Figure 3i, the intra-compartment interactions (the red curves) account for a higher proportion of the total contacts than the inter-compartment interactions (the grey curve). This is consistent with the typical patterns of intra- and intercompartment interactions that have been widely validated in prior research (2, 38).

From a functional perspective, compartment A is typically characterized by a greater abundance of active transcriptional activities, resulting in a higher number of active promoters and enhancers. Figure 3j and Supplementary Figure S2a provide a comparison of the active histone modification markers (H3K4me3 and H3K27ac as detected in bulk ChIP-Seq) in compartments A and B as detected by scCAFE. The analysis revealed a significantly higher enrichment of both markers in the compartment A loci. The enrichment of these markers were further elucidated in a stratified manner, where the mean number of ChIP-Seq peaks was plotted against the posterior probability of the compartment predictions (Figure 3k and Supplementary Figure S2b). The results indicate that our predicted compartment scores (posterior probabilities) can serve as indicators of the activity levels at the respective loci. This offers a potential reference for exploring the regulatory characteristics of genomic regions in single cells.

### scCAFE-annotated architectural features are predictors of cell identity

scHi-C is a powerful technique that enables the detection of cell-to-cell variability in the 3D genome across individual cells, as well as the heterogeneity across different cell types (18). Several existing studies have focused on projecting scHi-C data into a lower-dimensional space for the purpose of cell clustering (21, 23, 26, 39). However, the investigation of architectural features’ heterogeneity across different cell types has been relatively limited. In this study, we made use of the architectural features identified by scCAFE to explore how these features are related to the cell type identity. To this end, we generated four different sets of cell features for the hPFC dataset using scCAFE: the high-level latent features (i.e., the latent features generated by the multi-task VGAE), loop features, TLD features, and compartment features. Although scCAFE identifies all levels of architectural elements at 10 kb resolution, we pooled the predictions to 100 kb resolution so that the feature vectors of cells could fit into memory. Using our encoding scheme, all four sets of cell features were of the same dimensionality (Methods).

Here, we utilized unsupervised dimensionality reduction techniques to effectively visualize the cell embeddings. Additionally, we trained supervised machine learning models to assess the predictive capability of single-cell architectural features in determining cell types. Figure 4a display the UMAP visualizations of the four sets of features and the classification performance metrics obtained using these features. The 2D visualization (Figure 4a leftmost) of the latent features clearly demonstrates the separation of the seven different cell types in the hPFC dataset. The accuracy and macro-F1 score achieved were 0.99 and 0.93, respectively. The confusion matrix was visualized and displayed in Supplementary Figure S3. These results strongly indicate that the features extracted by scCAFE successfully captured the heterogeneity of the 3D genome organizations across cell types. In comparison, the single-cell loops and compartment features showed less capability in distinguishing between the cell types (Figure 4a and Supplementary Figure S3b,d). Among the four sets of features, the TLD features exhibited the least pre-dictive power in differentiating cell types. Despite this, the TLD features still demonstrated certain classification ability, as their metrics were higher than those of a random (dummy) classifier (accuracy = 0.40, macro-F1 = 0.08).

**Fig. 4.**
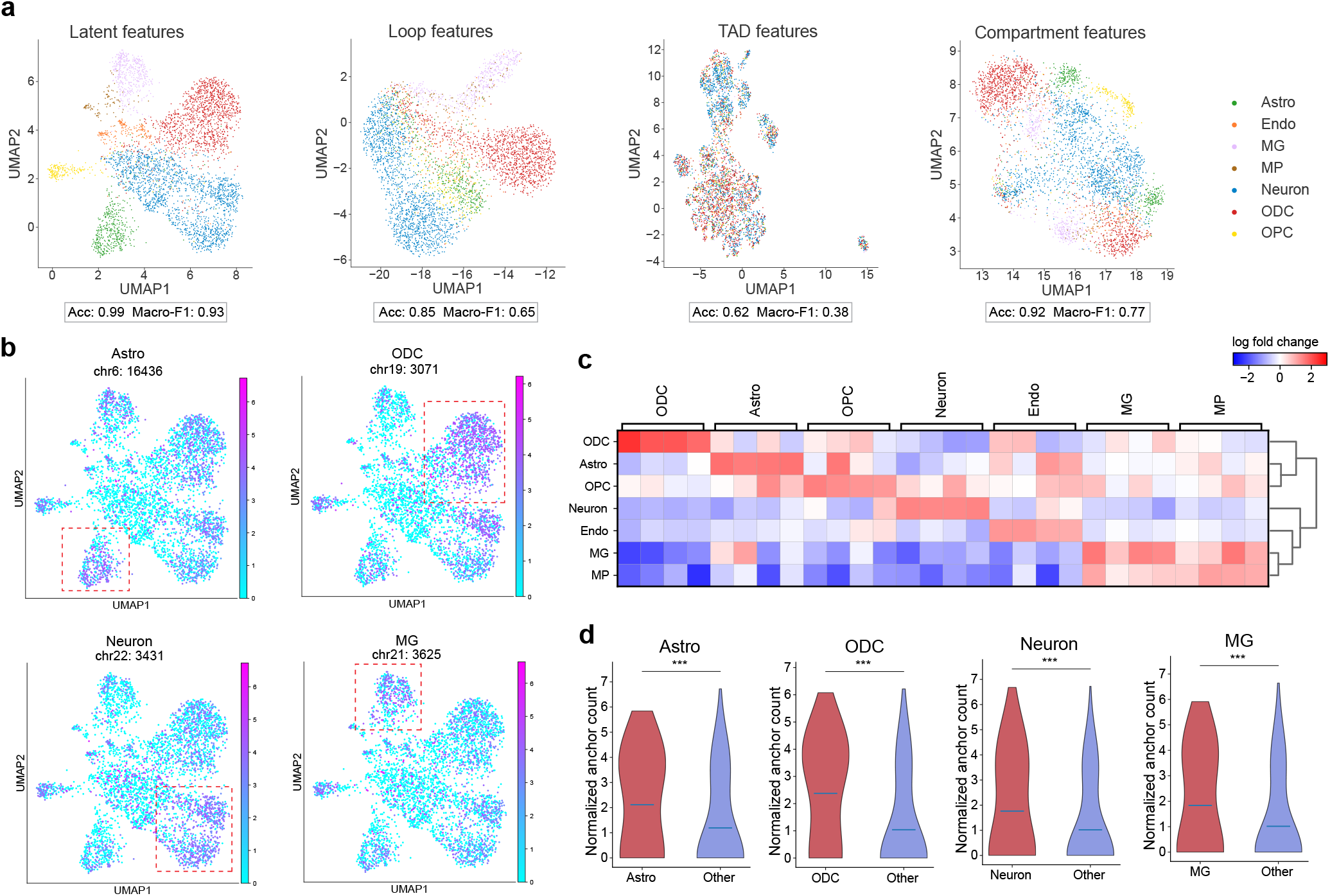
Predictive power of scCAFE-predicted single-cell architectural features in classifying cell types. **(a)** UMAP plots of scCAFE latent features, single-cell loops, TLDs, and compartments on the hPFC dataset. **(b)** “Marker loop anchors” identified by scCAFE in Astro, ODC, Neuron, and MG. The red square in each subplot denotes the enriched target cells. **(c)** Matrix plot of marker loop anchors. Each row represents a cell type, and each column corresponds to a loop anchor region in the genome. The color of each entry denotes the log fold change in the number of loops compared to other cell types. **(d)** Violin plots of the loops associated with the marker loop anchors in different cell types, comparing between the enriched cell type and other cell types. Three asterisks (***) denote p-value smaller than 0.001, Mann-Whitney U test.

The observations present significant insights into the variability of genome spatial organization. Notably, the highlevel latent features exhibited superior predictive power compared to all other feature sets, as anticipated. This finding aligns with the well-established principle of hierarchical structures within the 3D genome. In this context, different levels of single-cell architectural features can only reflect specific aspects of the 3D genome hierarchy. On the other hand, the cell-type heterogeneity of single-cell TLDs was not rich enough to support accurate cell type identification, indicating a more conservative role of domain boundaries across cell types. This is in line with multiple previous studies on the conservative nature of domain boundaries, which often serve as stable organizational units within the genome (33, 40, 41). Although it has been demonstrated in Figure 3a and d-g that the TLD boundaries were variable across single cells, this heterogeneity may only account for the dynamic shifting of domain boundaries, yet not significant enough for cell type identification.

Overall, the degree of variability across cells differs depending on the architectural features. In the next section, we examine single-cell loops as an example to further explore the connections between architectural features and cell identities.

### Marker anchors identification using single-cell loops predicted by scCAFE

The concept of the marker gene has been vitally important in single-cell analysis, as it allows for the identification and characterization of specific cell types (42–45). In this study, we generalized the concept of marker gene to provide a visualized and interpretable profile of how the loop anchors annotate cell types.

To achieve this, we adjusted the encoding scheme for singlecell loop anchors. The feature encoding scheme used in the former section was designed for a fair comparison between features, i.e., the feature dimensionality was identical across feature sets. This scheme encoded the features using 100 kb-resolution bins, ensuring the resulting vectors fit into memory. This was necessary despite the fact that the original features were called at a resolution of 10 kb. To gain a better understanding of the relationship between single-cell loops and cell types, we employed a more appropriate method to encode the single-cell loop anchors at a 10 kb resolution, where the loop anchors in each cell were input as a text document, and the tf-idf scheme was applied. In this way, the occurrence frequencies of loop anchors were well preserved in the resulting features.

By training seven SVM models (one for each cell type) using tf-idf features and cell type labels as inputs, we successfully developed a one-vs-rest multi-class classifier capable of accurately predicting cell identity. This classifier achieved an accuracy of 0.91 and a macro-F1 score of 0.78, indicating superior performance compared to the model using features from the previous section. By leveraging the weights learned by SVMs, we identified 50 candidate anchors for each cell type that exhibited the highest positive discriminative power. These candidate anchors were then subjected to further analysis using Scanpy (43) to recognize key anchors associated with Astro, MG, Neuron, and ODC cell types. Through our analysis, we discovered key anchors that were particularly informative for distinguishing between these cell types. These key anchors, as illustrated in Figure 4b-d, exhibited a substantial variation in the number of associated loops across different cell types. This finding suggests that the distribution and abundance of loop anchors can serve as valuable markers for characterizing and differentiating distinct cell populations.

In summary, our study highlights the potential of loop anchors as markers for cell type identification. This may illuminate future studies in single-cell 3D genomics by providing identification references for cell types solely from scHi-C data.

## Discussion

In this study, we proposed a computational framework scCAFE that can directly predict the architectural features of 3D genome at multiple levels without dense imputation of scHi-C contact maps. With this framework, we were enabled to detect chromatin loops with a unprecedentedly high accuracy, and simultaneously detect the TLDs and A/B compartments that are highly consistent with the biological patterns observed in previous research. As enabled by the single-cell architectural features detected, we characterized the heterogeneity of these features across cell types, and we discerned the predictive power of different levels of architectural elements in classifying cell identity. Another significant contribution of our study is the identification of a series of “marker loop anchors”. Although we only employed standard data science protocols for this purpose, these anchors have the potential to serve as valuable clues for cell type identification in downstream studies. This is a particularly interesting direction that might be worth exploring in the future, as previous studies have been depending on the simultaneous profiling of other omics data (32, 46, 47). This concept aligns with the differential interactions (DIs) proposed in (46), which refer to the contacts that are significantly more intense in one cell type compared to others. The key distinction between these two methodologies lies in the spatial dimensionality of interest: marker loop anchors generated in our study are 1D vectors, whereas DIs produce 2D paired loci as output. Conse-quently, the representation of the less sparse 1D marker loop anchors is likely to exhibit greater robustness and invariance against noise.

The core principle underlying scCAFE’s algorithm design also follows the multi-view nature of Hi-C data (28, 31). The embeddings for genomic bins in each single cell are extracted from the graph representation of contact maps, and the chromatin loops are also identified using the graph view. In contrast, the detection of higher-order single-cell architectural elements is based on the sequence view of the data, utilizing unsupervised learning techniques. Specifically, scCAFE employs multi-task learning to capture patterns in 3D genome architecture, thereby enhancing the representational capacity of the embeddings. Hence, scCAFE can directly predict the 3D architectural features without imputing the contact maps to a density equivalent to bulk data. This strategy differs from the previously common practice in scHi-C analysis, where sparse contact matrices are densified to identify architectural elements. By eliminating the requirement for dense imputation, this strategy provides a more efficient framework for architectural feature identification. Several recent studies have adopted this non-imputation approach to predict single-cell loops (28) and scTLDs (29, 30). Nonetheless, to our knowledge, scCAFE is the first comprehensive model capable of predicting all three levels of architectural features of the 3D genome without the need for imputation.

A potential future improvement of scCAFE could be the integration of epigenomics and transcriptomics data. This would allow for the illumination of the connections between epigenetic modifications and the 3D genome, as well as the role of 3D genome architecture in gene regulation. As previously mentioned, several single-cell assays have been developed to simultaneously probe 3D genomics and other omics data (32, 46, 47). However, the development of computational tools to analyze these data remains limited. By training a neural network to predict gene expression in single cells using scCAFE embeddings, it might be possible to apply interpretable deep learning techniques to identify the most critical sub-structures in the 3D genome organization responsible for specific gene expression states. This conceptual framework would be constructed on the strong foundation established by our scCAFE embeddings, which have demonstrated exceptional presentational capabilities.

## Methods

### Data representation and model architecture of sc-CAFE

The data representation method in scCAFE was inherited from our previous work scGSLoop (28). In particular, each cis-contact map of a chromosome within a cell was represented as a graph, where each genomic bin served as a node, and the contacts between bins were represented by edges connecting these nodes. Leveraging the multi-view nature of Hi-C data (28, 31), the node features were extracted from DNA sequence information by encoding k-mer and motif occurrences as vectors. Given a scHi-C dataset with *L* cells and *C* chromosomes per cell, the data can be mathematically represented as:

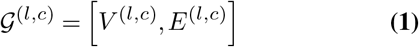

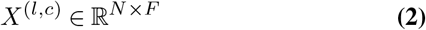

where *l* denotes the index of the cell in the dataset, and *c* denotes the *c*-th chromosome in the cell. *V* ^(*l,c*)^ and *E*^(*l,c*)^ represent the node set (genomic bins) and edge set (contacts) of the graph, respectively. Each row 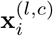 in *X*^(*l,c*)^ represents the feature vector of node *i* in the chromosome. All graphs were constructed using 10 kb-resolution contact maps.

In scCAFE, we employed a variational graph autoencoder (VGAE) (48)-like neural network to capture the high-level features hidden within the contact maps. We utilized a multiobjective loss function to optimize both the reconstruction of the original contact maps and the prediction of chromatin loops. Compared to scGSLoop (28), the auxiliary task of contact map reconstruction in scCAFE enabled the model to learn a more comprehensive representation of the chromatin organization, thereby allowing the prediction of multi-scale architectural features. Specifically, we treated the node embeddings in each graph as a sequence and applied unsupervised algorithms to detect higher-order organizational patterns in the genome, including TLDs and compartments.

Similar to scGSLoop, the scHi-C contact maps with low quality were enhanced using a k-nearest neighbors (KNN) algorithm before being input to the VGAE module. Initially, the cells were projected into a lower-dimensional space using scHiCTools (39). The contact matrices of each cell and its three nearest neighbors in this lower-dimensional space were averaged to create the input graphs for the VGAE.

### Multi-task VGAE for loop calling and node representation learning

In our approach, we employed a multi-task VGAE for node representation learning. We simultaneously optimized the VGAE for two related tasks: loop prediction and contact reconstruction. Both tasks involved edge-level predictions. For the loop prediction task, the model was trained to distinguish between positive looping node pairs, as specified in the reference loop list, and an equal number of negative non-looping pairs, which were sampled to match the count of the positive pairs. For the reconstruction task, the positive examples were the edges existing in the contact maps, and the negative samples were generated by randomly selecting non-existent edges within the contact maps. In both tasks, the proximity-aware negative sampling mechanism (28) was utilized to improve the model’s decision boundary and, consequently, enhance its learning capability.

The VGAE we adopted in this study was composed of a GraphSage (49) encoder and two edge-level dense decoders with identical architecture, each dedicated to a specific task. The forwarding rules can be formulated as follows:

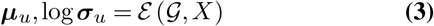

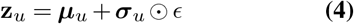

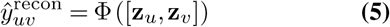

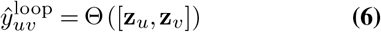

where ℰ, Φ, and Θ are the encoder, reconstruction decoder, and loop decoder, respectively. *ϵ* denotes a Gaussian distribution such that ϵ ∼ 𝒩 (0, **I**). *u* and *v* are indices of nodes in a chromosome graph, and [·, · ] represents the concatenation operation. In both tasks, the loss functions used were binary cross-entropy. The total loss was the sum of these binary cross-entropy losses and the KL loss.

The model trained in this multi-task manner can directly predict loops during the inference stage. Additionally, it can generate latent embeddings that can be utilized downstream for TLD and compartment detection.

### Detection of TLDs and compartments

The detection of single-cell TLDs and compartments was achieved using unsupervised learning methods. Here, the node embeddings within a chromosome graph, generated by the encoder ℰ, were treated as a linearly connected sequence corresponding to the genomic bins’ positions in the linear genome. For TLD pattern identification, we first reduced the dimensionality of embeddings using principal component analysis (PCA). We then applied hierarchical clustering with a connectivity constraint to determine the divisions of TLDs. This connectivity constraint ensured that the nodes within each resulting cluster were connected consecutively, preserving the sequential characteristics of the genomic bins in TLDs. The connectivity matrix *M* ^(*l,c*)^ ∈ ℝ^*N ×N*^ for each chromosome graph 𝒢^(*l,c*)^ is defined as:

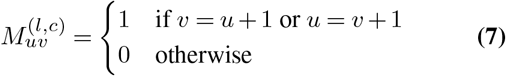

The number of clusters for each chromosome was calculated by dividing the chromosome length by 200 kb. In this way, a set of candidate TLD boundaries was identified. These candidate boundary bins were further categorized into two groups using KMeans clustering. The group with a higher number of CTCF motifs was selected as the final list of TLD annotations.

Single-cell compartments were identified using a hidden Markov model (HMM). By modeling the genomic bins in each chromosome as a sequence, we assumed that two hidden states underlie the observations, corresponding to compartments A and B. The HMM model was trained on the node embeddings of 100 chromosomes. The trained model was then used to predict the hidden states of nodes for each graph in the dataset. We subsequently adjusted the signs of the posterior probabilities based on GC content. Specifically, the state with higher GC content was assigned positive probability values, while the state with lower GC content was assigned negative probability values. This sign flipping scheme was consistent with the practices in previous studies (6, 50, 51) and was also adopted in existing Hi-C analysis libraries (52).

Both TLDs and compartments were detected at the resolution of 10 kb. The resolution of the predictions was then coarsened to meet the requirements for downstream analysis as needed.

### Consensus architectural feature prediction

scCAFE also provides methods for the consensus architectural features of a cell population, such as cell type or subtype. For consensus loop detection, we followed the simple aggregation method used in (28), which calculates the average loop probability across the cell population.

To obtain the consensus TLD predictions, a more complex method is required. First, a co-association matrix was created by recording the number of cells in which each pair of nodes was assigned to the same cluster. This co-association matrix was then divided by the total number of cells, resulting in an adjacency matrix. The adjacency matrix was utilized to obtain the spectral embedding. Finally, hierarchical clustering was applied to the spectral embedding, effectively dividing the nodes within a chromosome into multiple TLD regions. For the consensus predictions of compartments, we utilized the GC-flipped posterior probabilities of the loci and calculated the average across all cells within the population.

### Embedding and classification using latent and architectural features

We examined the predictive power of different sets of features (including the latent high-level features and single-cell architectural features) in determining cell identity in the human prefrontal cortext (hPFC) dataset. This was achieved by visualizing the 2D embeddings of cells and conducting supervised classification on the cells.

To create the latent high-level features that can fit into memory, we first employed PCA to reduce the dimensionality of the output from the encoder ℰ of scCAFE. Only the first principal component was retained in our study, resulting in each chromosome graph having a single feature vector. This chromosome feature vector was then pooled by summing up every 10 dimensions, so that the final dimensionality was consistent with the number of bins in the 100 kb-resolution contact map. Architectural features were also aligned to 100 kb resolution. Loop features were generated by taking the numbers of loops associated with each 100 kb anchor. The presence of TLD boundaries in 100 kb bins was converted into a binary vector, which was subsequently used as the TLD features. Compartment features were the posterior probabilities from HMM coarsened to 100 kb resolution by taking the average of every 10 entries. The latent high-level features, loop features, TLD features, and compartment features were all at 100 kb resolution, ensuring they have the same dimensionality.

Each set of features was embedded into a 2D space using a combination of PCA and UMAP. PCA initially reduced the dimensionality, and then UMAP projected the lower-dimensional representations into 2D for visualization. Supervised machine learning method was then used to quantify the discriminative power of the features. In particular, seven linear support vector machines (SVMs) with L1 regularization were trained for each set of features to classify the cells into seven types. The training set for each feature set was composed of 75% of the cells, and the left cells were used for testing the model’s performance.

### Discovery of marker loop anchors

Combining traditional machine learning techniques and statistical methods, we identified marker loop anchors for the seven cell types in the hPFC dataset. Different from the encoding scheme described in the previous section, we took the loop anchors of 10 kb resolution, and subsequently encoded them utilizing the tf-idf scheme. A total of seven linear SVM models were trained using these features. Each model was specifically optimized to classify cells of a particular cell type from the remaining cells in the dataset. The learned weights of each SVM were then considered as the contributions of the features (anchor loci) to the classification outcomes. We selected the top 50 contributing features with the highest positive weights for each cell type. The de-duplicated union of these loci, identified across all cell types, constitutes the pool of candidate markers. Scanpy (43) was utilized to further identify marker loop anchors for each cell type by performing statistical tests across cell types. Among the loci in the candidate marker pool, it identified those that are associated with a significantly higher number of chromatin loops compared to other cell types. This analysis was based on the assumption that the distribution of the number of associated loops is similar to that of the expression volume of RNA, which was modeled in Scanpy. We selected the default method for statistical test (t-test) in Scanpy, and the correction method was Benjamini-Hochberg.

The classification performance using these tf-idf features was also evaluated and is reported in this manuscript.

### Metrics for performance evaluation

In this study, we employed F1 score and precision-recall curve (PR-curve) to evaluate the models’ performance in terms of loop calling. The F1 score is a metric that effectively combines precision and recall into a single value, striking a balance between these two measures. However, calculating the F1 score requires choosing a cutoff point to convert the probabilities generated by neural networks into binary predictions. Consequently, this metric may be less suitable when an extensive evaluation across all thresholds is required. To overcome this issue, we additionally assessed the performance of the models using PR-curve for a comprehensive evaluation.

The metrics were calculated based on the entries within the genomic distances ranging from 100 kb to 1 Mb on the contact maps, which corresponded to the regions where the predictions were made. It is worth noting that, different from the slack F1 score used in (28) and (22), here we adopted the original definition of precision and recall to calculate F1 score and plot the PR-curve.

The metrics for evaluating the cell type classification results include the F1 score, accuracy, and the confusion matrix. Given the multiclass classification setting, we calculated the macro-F1 scores to assess performance. Accuracy measures the overall correctness of the classification, while the confusion matrix provides a visual representation of the algorithm’s detailed performance.

### Architectural features called from bulk Hi-C data

We used the same reference loop lists for the mESC dataset and the hPFC dataset as the ones used in SnapHiC (22) and scGSLoop (28). The reference loops were called from bulk Hi-C data and were used as labels for training the model and evaluating the model’s loop predictions.

The insulation scores of genomic bins were calculated from bulk Hi-C data using the `insulation` function in FANC (52). These scores were obtained with window sizes of 1 Mb, 1.5 Mb, 2 Mb, 2.5 Mb, 3 Mb, 3.5 Mb, and 4 Mb.

Per-chromosomal normalization and log transformation were applied to the scores.

To generate the bulk A/B compartment annotations, we applied eigendecomposition on the Pearson correlation matrix of each contact map at 100 kb resolution. The resulting first eigenvalues were then flipped using GC content such that the entries in A compartment have positive values, while entries in the B compartment have negative values. This A/B compartment calling process was performed using the `compartments` function in FAN-C package.

### Aggregate analysis

The aggregate analyses of loops, TLDs, and compartments were also carried out using FAN-C (52). For the loop aggregate analysis, we selected the top 5,000 loops with the highest predicted probabilities, and aggregated the looping pixel and the surrounding pixels. The TLD aggregation was conducted following the method in (14). The contacts within TLDs and their surrounding regions were cropped into square matrices and re-scaled to a uniform size. These matrices were subsequently piled up to produce the final aggregate figure. We employed a saddle plot to create the aggregate profile for compartments. Each entry on the plot was the average O/E contact between the regions represented by the row and those represented by the corresponding column.

All aggregate analyses were conducted on the same bulk Hi-C data, utilizing annotations derived from the single-cell dataset (including both single-cell annotations and consensus annotations). This approach guaranteed the consistency of data sources for aggregation and ensured the comparability of the aggregate results. For TLD aggregation, only TLDs of size larger than 100 kb and smaller than 1 Mb were considered.

## Code availability

The code of scGSLoop is open-source and publicly available on GitHub at https://github.com/fzbio/scCAFE.

## Data availability

The data used in this study are all publicly available. No new data were generated in this study. The mESC scHi-C data, initially generated by Nagano et al. (12), are accessible via the Gene Expression Omnibus (GEO) with the accession number GSE94489. Raw data for the hPFC dataset (32) are available on GEO under accession number GSE130711. For the two datasets, we used the processed contact maps which are available at https://github.com/tanaylab/schic2 and https://salkinstitute.app.box.com/s/fp63a4j36m5k255dhje3zcj5kfuzkyj1/folder/82403061106, respectively. CTCF motif annotations for mm10 and hg19 were collected and processed from (53). These annotations were generated by scanning the whole genomes for CTCF motifs with FIMO (54).

## ACKNOWLEDGEMENTS

This research was substantially sponsored by the research project (Grant No. 32170654) supported by the National Natural Science Foundation of China and was substantially supported by the Shenzhen Research Institute, City University of Hong Kong. The work described in this paper was substantially supported by the grant from the Research Grants Council of the Hong Kong Special Administrative Region [CityU 11203723]. The work described in this paper was partially supported by the grants from City University of Hong Kong (2021SIRG036, CityU 9667265, CityU 11203221) and Innovation and Technology Commission (ITB/FBL/9037/22/S).

