## Supplementary Figures for "Unveiling multi-scale architectural features in single-cell Hi-C data using scCAFE"

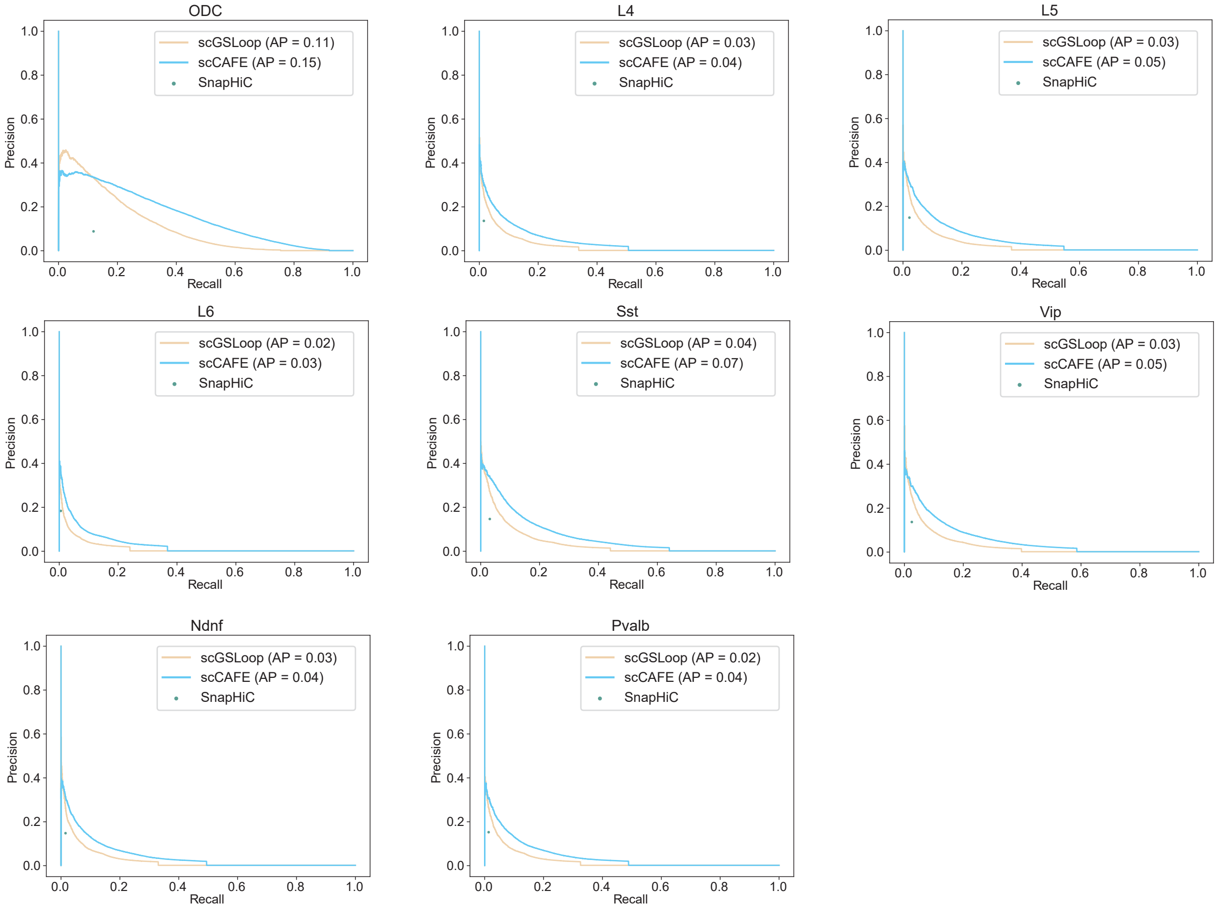


**Figure S1.** Precision-recall plots of the consensus loops in ODC, L4, L5, L6, Sst, Vip, Ndnf, Pvalb cells predicted by SnapHiC, scGSLoop, and scCAFE.

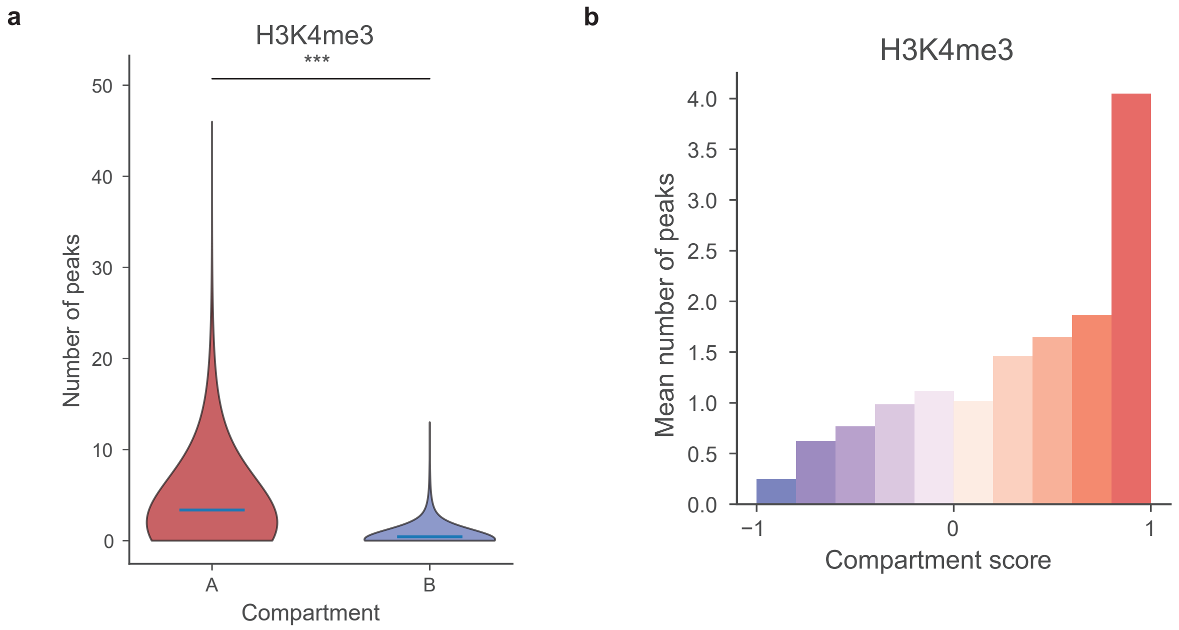


**Figure S2.** **(a)** Numbers of H3K4me3 ChIP-Seq peaks in compartment A and B. **(b)** Numbers of H3K4me3 peaks at different scCAFE compartment scores.

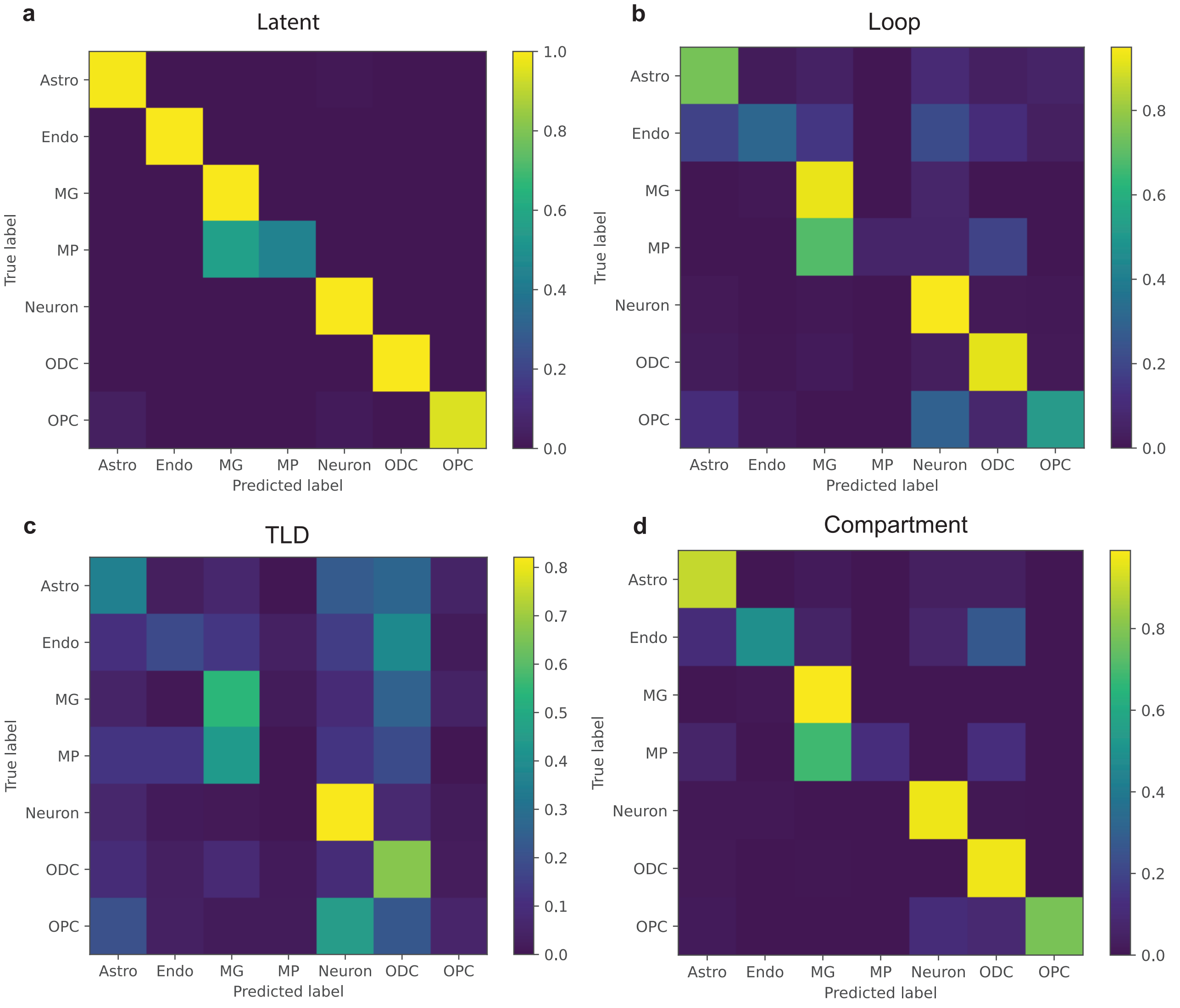


**Figure S3.** Confusion matrices of different sets of architectural features. **(a)** scCAFE latent features. **(b)** Single-cell loop features. **(c)** Single-cell TLD features. **(d)** Single-cell compartment features.
